# Sharp cell type boundaries emerge from coordinated morphogen signaling

**DOI:** 10.64898/2026.04.02.716142

**Authors:** Ruiqi Li, Yiqun Jiang, Sarah Platt, Tianchi Xin, Ryan Driskell, Kevin A. Peterson, Sarah Van, Hainan Lam, Shagun Lukkad, Eva-LaRue Barber, Chae Ho Lim, M. Mark Taketo, Yuval Kluger, Peggy Myung

## Abstract

Classic models of the French flag problem depict sharp cell-type boundaries emerging from threshold responses to morphogen gradients. How discrete cell-type boundaries arise from morphogen signals that vary continuously across developing tissues is not completely understood. We use hair follicle dermal condensate formation to study a sharp developmental transition in which proliferative progenitors undergo cell-cycle exit concurrent with molecular differentiation. Using genetic and genomic approaches, we show that Wnt and Hedgehog signaling interact to coordinate the timing of these two processes. We identify a division of labor between the pathways: Wnt signaling promotes cell-cycle exit by regulating chromatin binding of the Hedgehog mediator GLI3, while Hedgehog signaling induces differentiation genes in a Wnt-dependent manner and simultaneously elevates Wnt activity. When Wnt and Hedgehog activities are temporally aligned, differentiation and cell-cycle exit occur within the same developmental window, restricting both the duration and abundance of intermediate states and producing a sharp cell-type boundary. When these signals are misaligned, intermediate states persist and expand, producing fuzzy boundaries. These findings reveal a mechanism in which interacting morphogen signals regulate the duration and abundance of intermediate states during a developmental transition, thereby controlling how continuous cell-state progression is translated into discrete tissue patterning.

## Introduction

Developing tissues must position distinct cell types across space and time to assemble functional organs. Lewis Wolpert framed this challenge as the “French flag” problem, in which discrete cell types emerge at defined positions along a morphogen gradient (Fig. 1A) (Wolpert 1969). In this framework, graded morphogen signals are interpreted through concentration thresholds that delineate spatial domains of cell fate (Driever and Nusslein-Volhard 1988; Roelink et al. 1995; Benzinger and Briscoe 2025). Yet, a fundamental question remains unresolved: how sharp cell-type boundaries emerge *in vivo* when the underlying signals and cell states vary continuously across a tissue. In developing systems, boundaries do not arise from instantaneous fate switches. Instead, cells progress through dynamic transitions and pass through intermediate states before committing to a new identity. In many mammalian tissues, these transitions unfold while cells continue to proliferate, raising the question of how ongoing cell-cycle dynamics are coordinated with graded morphogen inputs. Thus, boundary sharpness may reflect not only where thresholds are crossed, but how rapidly cells traverse a shared developmental progression.

**Fig. 1:**
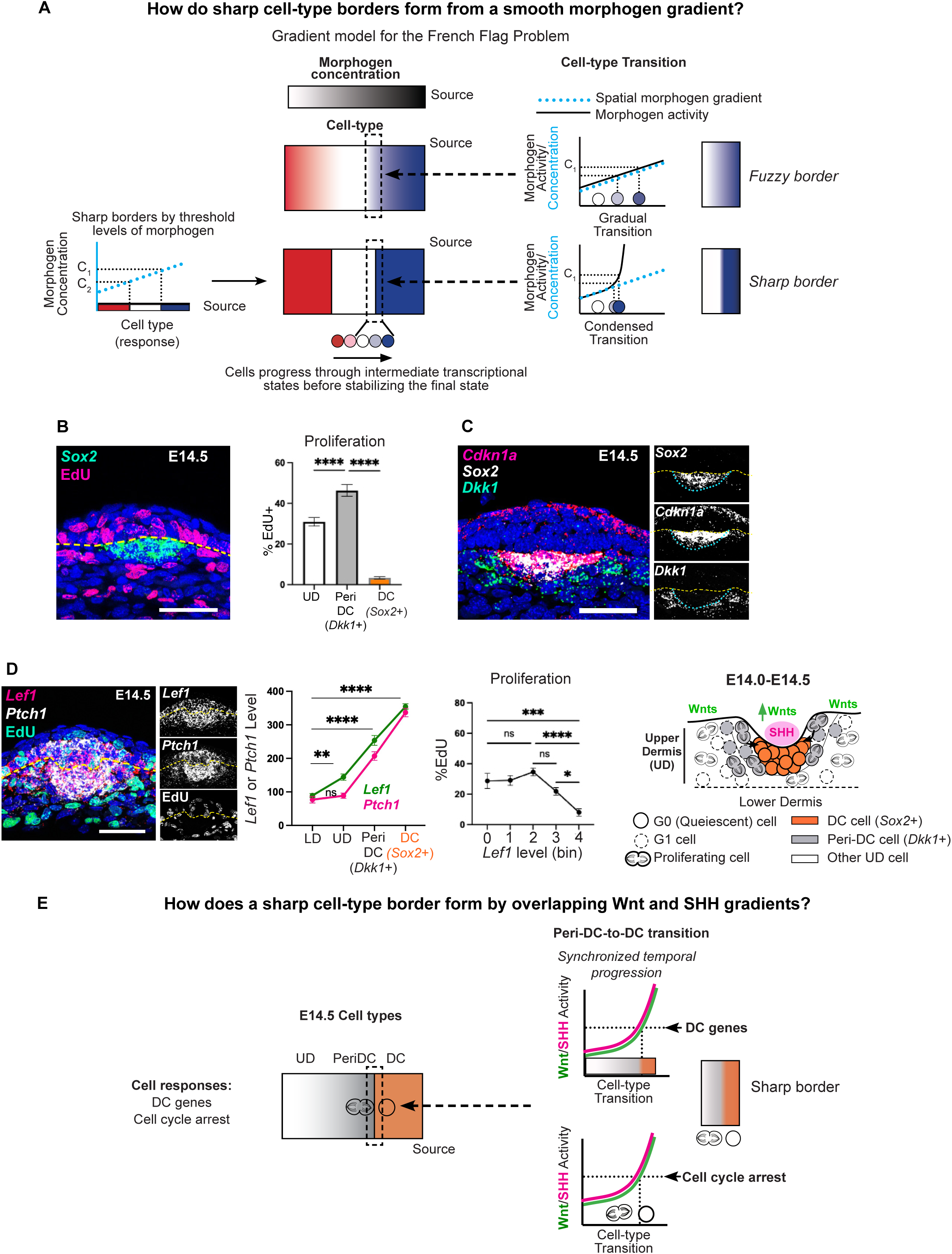
Cell-cycle exit and molecular differentiation are tightly coupled during dermal condensate formation. (A) French flag problem, illustrating threshold-based morphogen patterning (left) and alternative dynamic transitions across a fixed morphogen gradient (right) with either linear responses or non-linear transitions across morphogen levels. (B) FISH of E14.5 skin showing *Sox2*+ DCs lacking EdU incorporation; quantification of %EdU by cell-type (UD, upper interfollicular dermis) (n=10). (C) FISH showing peri-DC *Dkk1*^+^ cells adjacent to *Sox2*^+^ DCs with *Cdkn1a* enrichment at the DC boundary (n=10). (D) E14.5 skin stained for *Lef1* and *Ptch1*; transcript levels quantified by spatial regions (LD, lower dermis) or %EdU+ by *Lef1* bin (n=5). (E) Schematic summarizing dermal cell states relative to spatial position and Wnt/SHH activity. DCs arise in regions of overlapping Wnt and SHH activity, where cell-cycle exit coincides with DC gene expression occur. Data as mean± SEM; one-way ANOVA; ns, not significant. Scale bars=50 µm.

Addressing the temporal dimension of the French flag problem has been challenging, as most approaches to studying cell fate transitions emphasize cell origins or endpoints, rather than intermediate regulatory programs, or can visualize cell behaviors without molecular resolution (Rompolas et al. 2012; Ouspenskaia et al. 2016; Mok et al. 2019; Morita et al. 2021). We address this problem in a mammalian system by examining hair follicle dermal condensate formation, a tractable developmental transition in which the principal morphogen signals are known and cell-state progression occurs over a narrow temporal window (Chuong et al. 1996; Glover et al. 2017; Gupta et al. 2019).

Dermal condensates (DCs) are clusters of quiescent dermal cells that form beneath epithelial hair follicle placodes and are essential for hair follicle development (Chase et al. 1951; Hardy 1992; Millar 2002). At E13.5, prior to DC formation, Wnt signaling is broadly activated across the upper dermis in response to epidermal Wnt ligands, forming a gradient that is highest in cells closest to the epidermis (Noramly et al. 1999; Chen et al. 2012; Gupta et al. 2019). Wnt activity is required but not sufficient for DC formation. Upon placode formation, placode epithelial cells secrete Sonic Hedgehog (SHH) and additional Wnt ligands, resulting in localized co-activation of Wnt and SHH in underlying dermal cells (St-Jacques et al. 1998; Chiang et al. 1999; Gritli-Linde et al. 2007; Woo et al. 2012). Together, these signals are necessary and sufficient for DC formation, yet how Wnt and SHH signaling interact to drive the pre-DC-to-DC transition remains unclear, and DCs appear to emerge abruptly with few detectable intermediates (Glover et al. 2017; Gupta et al. 2019).

Previous work has examined how tissue growth and proliferation affect morphogen gradients by shaping field size, ligand distribution, or the scaling of positional information (Lander et al. 2002; Gregor et al. 2007; Kicheva et al. 2012). In these models, proliferation is typically considered in terms of how it modifies the signaling environment cells interpret. However, many mammalian tissues undergo progressive differentiation while cells continue to divide, introducing an additional constraint, as proliferative expansion of intermediate states may influence the precision of pattern formation. Here, we examine how interacting morphogen signals coordinate cell-cycle exit with differentiation to ensure precise spatial patterning in proliferative tissues.

Combining *in vivo* genetic perturbations, chromatin profiling, and a single-cell computational framework that disentangles concurrent biological programs, we capture intermediate states that are normally difficult to detect and determine how their timing and spatial extent are regulated by Wnt and SHH signaling *in vivo* (Qu et al. 2024). We show that Wnt and SHH differentially regulate cell-cycle exit and differentiation, and that their temporal coordination determines whether intermediate states are compressed or expanded, thereby controlling boundary sharpness. Mechanistically, we identify cross-regulation of transcription factor chromatin occupancy as a key mechanism linking signaling dynamics to cell-cycle exit.

## Results

### Cell-cycle exit and molecular differentiation are temporally coordinated during dermal condensate formation

Classic models of the French flag problem posit that distinct cell states are specified by threshold levels of morphogen concentration that vary with distance from a polarized source, predicting that sharp cell-type boundaries arise from spatial differences in morphogen signaling (Fig. 1A). To examine how this model applies to hair follicle dermal condensate (DC) formation, we asked how dermal cell types are spatially organized before and during DC formation. Prior to DC formation, dermal cells are proliferative and lack DC-specific markers (Supplemental Fig. 1A). Upon DC formation, DC cells express characteristic differentiation markers, including *Sox2*, *Sox18*, and *Gal*, and undergo cell-cycle exit, as indicated by minimal EdU nucleotide incorporation, while neighboring upper dermal cells that do not express DC genes remain highly proliferative. (Fig. 1B,C; Supplemental Fig. S1B,C).

Using *in vivo* labeling, we previously showed that DC cells arise from highly proliferative progenitors located in the upper dermis at E13.5 (Supplemental Fig. S1D) (Gupta et al. 2019; Qu et al. 2022). As DC formation initiates, proliferating cells that give rise to DCs become localized to a region surrounding the nascent condensate (peri-DC region) where they express *Dkk1* (Fig. 1C and Supplemental Fig. S1C). During the pre-DC–to–DC transition, *Dkk1*⁺ peri-DC cells exit the cell cycle while upregulating DC differentiation genes (Qu et al. 2022). Accordingly, DC marker expression overlapped with the *in vivo* G1/G0 Fucci2 reporter and with expression of the cell-cycle inhibitor *Cdkn1a* (*p21*), with only rare cells expressed *Cdkn1a* in the absence of *Sox2* (Fig. 1C; Supplemental Fig. S1E) (Biggs et al. 2018; Gupta et al. 2019). These data indicate that cell-cycle exit and DC differentiation occur concurrently during DC formation.

Given this temporal coincidence, we next asked how these two processes relate to Wnt and Sonic Hedgehog (SHH) signaling during DC formation. At E13.5, prior to SHH expression, Wnt signaling is active in the upper dermis (Supplemental Fig. S1A) but is not sufficient to induce DC gene expression (Zhang et al. 2009; Chen et al. 2012; Gupta et al. 2019). Using *Lef1* as a canonical readout of Wnt activity, which is concordant with expression of multiple Wnt target genes (*Axin2*, *Wif1*, *Tcf7*) (Supplemental Fig. S1F) (Gupta et al. 2019; Qu et al. 2022), we examined the relationship between Wnt signaling and proliferation. At E13.5, *Lef1* expression was highest in the most superficial layer of upper dermis and decreased across deeper dermal cell layers, yet this spatial pattern did not correlate with EdU incorporation, nor with DC gene expression (Supplemental Fig. 1A).

By E14.5, SHH expression in placode cells resulted in SHH activation in the underlying dermis. At this stage, dermal SHH activity (assessed by *Ptch1*) covaried with Wnt signaling (*Lef1*) across dermal populations, with significantly higher *Lef1* levels detected compared to E13.5 (Fig. 1D). The highest levels of both signals were observed in DC cells, intermediate levels in peri-DC (*Dkk1*⁺) progenitors, and progressively lower levels in surrounding upper dermal cells. In contrast to E13.5, elevated Wnt activity at E14.5 correlated with reduced proliferation, with the lowest EdU incorporation observed in DC cells (Fig. 1D,E). Thus, quiescence and molecular differentiation coincide spatially and temporally within a narrow domain characterized by elevated Wnt and SHH signaling.

### High Wnt activity induces cell-cycle exit independently of SHH signaling

To examine how Wnt and SHH signaling contribute to the pre-DC–to-DC transition, we perturbed each pathway prior to DC formation. Because both pathways are required for DC differentiation, loss-of-function approaches preclude analysis of their individual roles at this stage. As DC cells ultimately exhibit the highest levels of both Wnt and SHH activity, we asked whether these signals contribute to distinct components of the transition or act through a shared mechanism. We therefore elevated Wnt signaling before DC formation by expressing constitutively activated β-catenin (*Axin2CreER;βcat^fl/EX3^*, hereafter Actβcat; Fig. 2A, tamoxifen at E11.5).

**Fig. 2:**
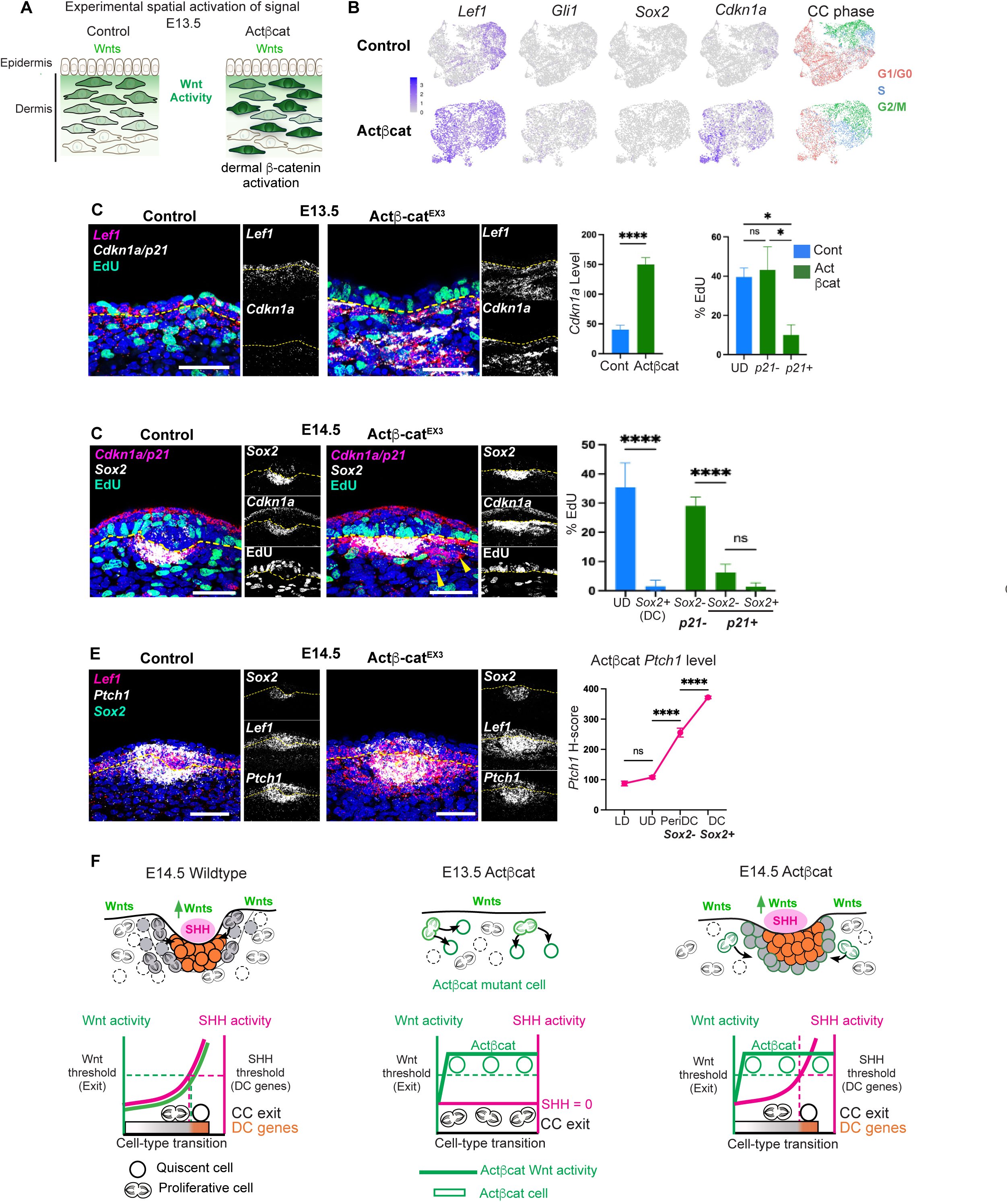
High Wnt activity induces cell-cycle exit independently of SHH signaling. (A) Experimental design for constitutive dermal β-catenin activation before DC formation (tamoxifen E11.5). (B) UMAPs of E13.5 Actβcat mutant and control dermal scRNA-seq data colored by indicated genes or cell-cycle phase (n=2). (C) FISH for *Lef1*, *Cdkn1a*, EdU in E13.5 skin with quantification of *Cdkn1a* transcripts (H-score) and %EdU of *Cdkn1a* (*p21*)^+^ or *p21*^-^UD cells (n=3). (D) E14.5 FISH for *Sox2*, *Cdkn1a*, EdU with quantification of %EdU^+^ cells in control or Actβcat *Cdkn1a*^+^ populations (n=5). (E) *Lef1*, *Ptch1*, *Sox2* FISH in E14.5 skin with quantification of *Ptch1* levels in Actβcat by cell type. (F) Model illustrating that elevated Wnt activity induces cell-cycle exit at E13.5 independently of SHH. DC molecular differentiation is restricted to cells with both high SHH and high Wnt activity. Scale bars=50 µm.

To assess how increased Wnt activity alters dermal transcriptional states prior to DC differentiation, we performed single-cell RNA sequencing of E13.5 Actβcat and control dermis. Dermal cells were filtered using established markers (e.g., *Col1a1*, *PDGFRα*). Consistent with prior work, Actβcat mutant dermal cells showed increased expression of Wnt targets (e.g., *Lef1*, *Axin2*, *Tcf7*), but not DC markers (e.g., *Sox2*) or SHH targets (e.g., *Gli1*) (Fig. 2B). Notably, *Cdkn1a*, a cell-cycle inhibitor normally expressed in DC cells at E14.5, was among the most significantly upregulated genes in E13.5 Actβcat dermis (Fig. 2B; Supplemental Table S1). Quantitative FISH confirmed elevated *Lef1* expression relative to controls and showed that Wnt target levels did not exceed those observed in wildtype DC cells, indicating that activated β-catenin increases Wnt signaling within physiological levels (Fig. 2C; Supplemental Fig. S2A). Constitutive Wnt activation was sufficient to induce *Cdkn1a* (*p21*) expression at E13.5 in the absence of SHH activation, and *Cdkn1a*+ cells exhibited reduced EdU incorporation (Fig. 2C).

By E14.5, DCs were present in both control and Actβcat embryos but were enlarged in mutants (Supplemental Fig. S2B). In controls, *Cdkn1a* expression was largely restricted to *Sox2*+ DC cells, whereas in Actβcat mutants, it extended into surrounding *Sox2*-upper dermal cells (Fig. 2D). Both *Sox2*+ and *Sox2*− cells expressing high *Cdkn1a* showed low EdU incorporation (Fig. 2D). *Sox2*+ cells in Actβcat mutants were closest to placodes and expressed the highest *Ptch1* levels, whereas surrounding mutant *Cdkn1a*+*Sox2*-cells expressed lower *Ptch1* levels, indicating that DC gene expression requires SHH signaling above a defined threshold (Fig. 2E). Thus, elevated Wnt signaling is sufficient to induce cell-cycle exit in the absence of SHH, whereas DC differentiation requires additional SHH-dependent input (Fig. 2F).

### SHH signaling cell-autonomously elevates Wnt activity coincident with cell-cycle exit

Our results show that elevated Wnt activity induces exit without DC differentiation. We next asked whether SHH activation could initiate DC gene expression independently of cell-cycle exit. To test this, we expressed constitutively activated Smoothened (*Axin2CreER;RosaLSL-Smom2YFP*, hereafter ActSMO) in the dermis prior to DC formation when Wnt signaling levels are low (tamoxifen E11.5; Fig. 3A). Whole-mount analysis of E14.5 skin showed that SHH activation in Wnt-active dermal cells induced large dermal clusters expressing DC genes in the upper dermis (Fig. 3B). These clusters formed autonomously without placodes, indicating that DC gene expression can occur without placode-derived Wnt ligands (Qu et al. 2022). Many cells in the outer layers of Sox2+ clusters remained proliferative (%Sox2*+*/EdU+), demonstrating that DC gene expression can occur independently of cell-cycle arrest. By contrast, Sox2+ cells in cluster centers lacked EdU incorporation similar to quiescent DC cells in wildtype embryos.

**Fig. 3:**
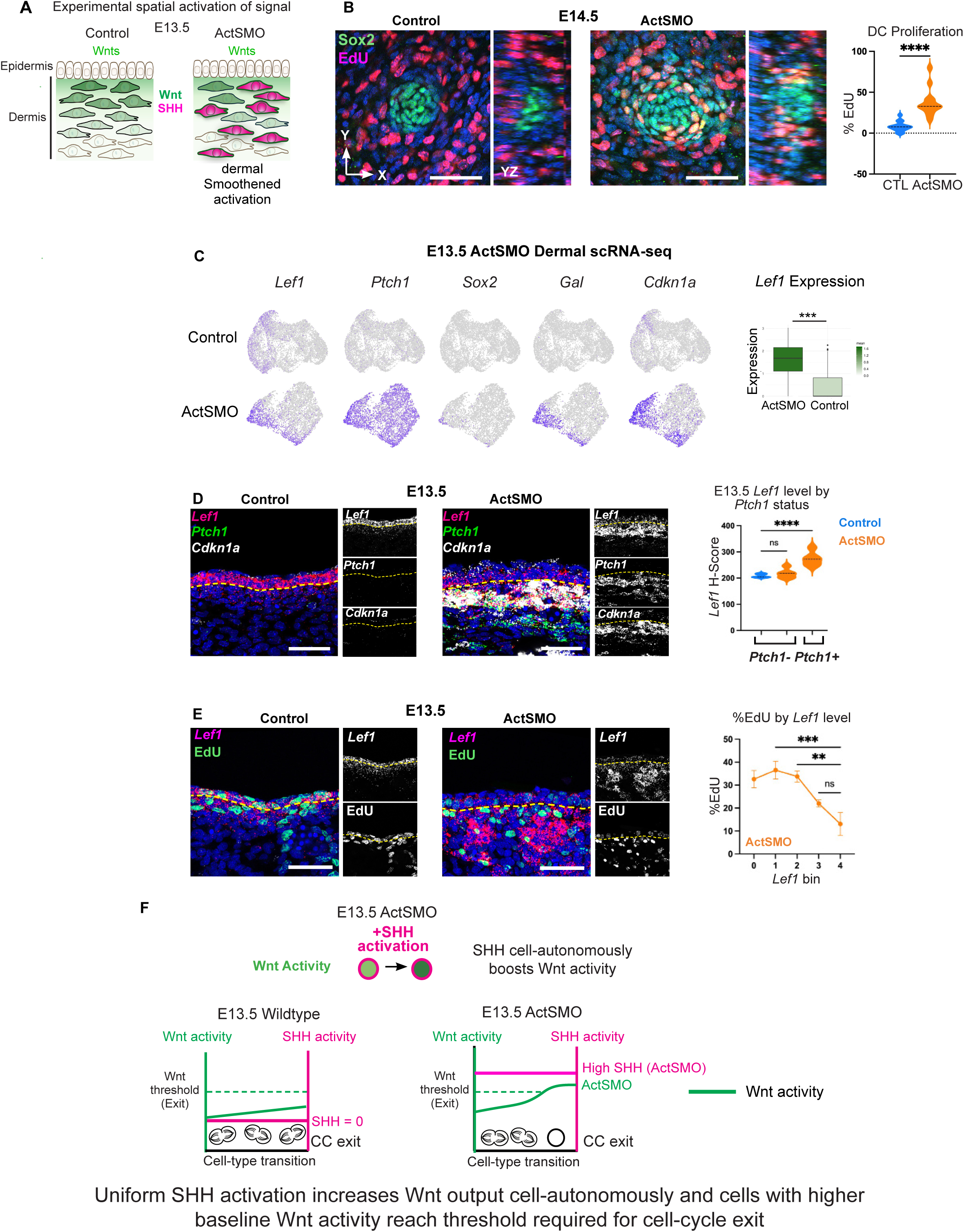
SHH signaling cell-autonomously elevates Wnt activity in association with cell-cycle exit. (A) Experimental design or constitutive dermal Smoothened activation prior to DC (tamoxifen E11.5). (B) Whole-mount E14.5 control and ActSMO skin stained for EdU and Sox2; quantification of %EdU+ in Sox2+ cells (n=8). (C) UMAPs of E13.5 control and ActSMO dermal scRNA-seq data colored by indicated genes; right, *Lef1* expression in UD cells. (D) E13.5 skin stained for *Lef1*, *Ptch1* and *Cdkn1a* with *Lef1* levels by *Ptch1* status (n=8). (E) E13.5 skin showing increased *Lef1* expression and reduced EdU incorporation in ActSMO dermal cells. (F) Model illustrating SHH-dependent elevation of Wnt signaling coincident with cell-cycle exit. Scale bars, 50 µm.

Given the observed quiescent central Sox2+ dermal cells (Fig. 3B), we examined whether similar molecular changes occurred in the E13.5 Actβcat mutant. scRNA-seq analysis of E13.5 ActSMO and control dermis showed uniformly high *Ptch1* expression in ActSMO cells (Fig. 3C). *Lef1* expression remained enriched in upper dermal (UD) cells but was significantly elevated compared with controls. FISH confirmed increased *Lef1* expression in ActSMO dermis despite the absence of placodes (Fig. 3D). Importantly, only *Ptch1*+ (recombined) cells showed elevated *Lef1* expression, while neighboring *Ptch1*-cells (non-recombined) retained normal *Lef1* levels, indicating a cell-autonomous effect. ActSMO cells with high *Lef1* also exhibited elevated *Cdkn1a* expression and reduced EdU incorporation (Fig. 3D,E). Together, these results show that SHH activation elevates Wnt signaling cell-autonomously and is associated with induction of cell-cycle exit, independent of placode Wnt ligands (Fig. 3F).

### GeneTrajectory resolves transcriptional programs associated with cell-cycle exit

In the ActSMO condition, DC genes are expressed in proliferating cells (Fig. 3B), indicating that exit and DC gene expression can be temporally uncoupled. To resolve the gene programs underlying these processes, we applied GeneTrajectory (GT), a computational approach that separates co-occurring gene programs from whole transcriptome scRNA-seq data (Qu et al. 2024).

Applying GT to E14.5 wildtype dermal scRNA-seq data identified three gene trajectories, corresponding to proliferation (S/G2/M), DC differentiation and lower dermal differentiation (Fig. 4A). The DC trajectory was partitioned into stages using gene bins and visualized by gene-bin scores (Fig. 4A; Supplemental Fig. S3A). *Lef1* and *Ptch1* expression covaried across DC trajectory stages and peaked at the terminal stage 6 (Fig. 4B; Supplemental Fig. S3B). Canonical DC genes, including *Gal*, *Sox18* and *Sox2,* were similarly enriched at stage 6 (Fig. 4C). Stage 6 cells were largely quiescent with a G1 fraction near 1.0 and elevated *Cdkn1a* expression, indicating that exit coincides with and molecular DC differentiation at the terminus of the trajectory.

**Fig. 4:**
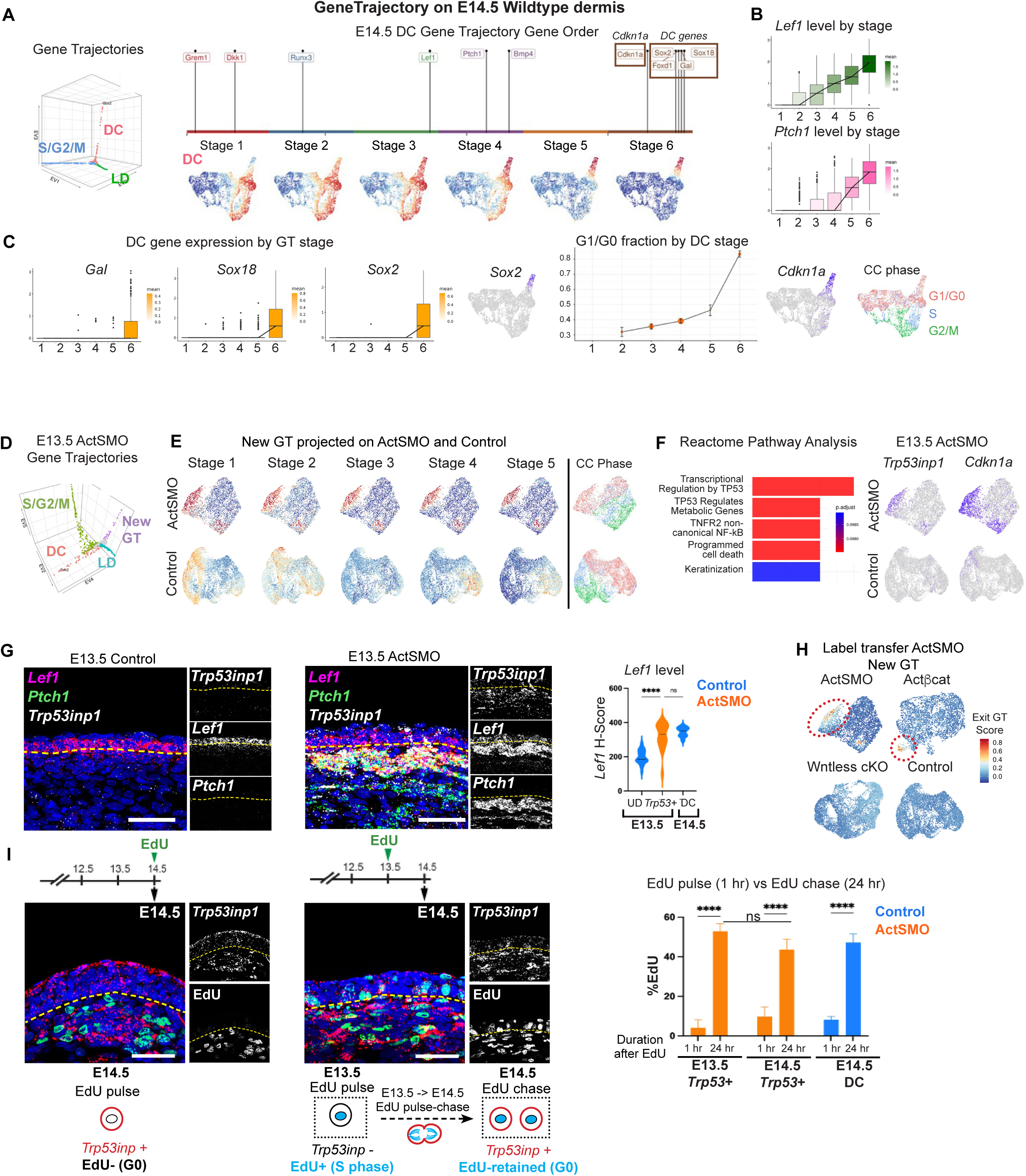
GeneTrajectory defines transcriptional changes accompanying cell-cycle exit. (A) GeneTrajectory diffusion map and gene order of E14.5 wildtype DC trajectory with dermal UMAPs colored by stage. (B) *Lef1* and *Ptch1* expression by DC trajectory stage. (C) DC gene expression and fraction of G1/G0 cells by DC trajectory stage with UMAPs of *Cdkn1a* expression and cell-cycle (CC) phase. (D) GeneTrajectory diffusion map of E13.5 ActSMO dermal cells. (E) Control and ActSMO dermal UMAPs colored by mutant trajectory stages or cell-cycle phase. (F) Pathway analysis of genes from new mutant trajectory with UMAPs colored by indicated genes. (G) FISH of E13.5 control and ActSMO skin for indicated genes with quantification of *Lef1* in upper dermal (UD), DC cells and ActSMO *Trp53inp1*+ cells (n=3). (H) Label transfer of the ActSMO new trajectory onto indicated E13.5 datasets showing cells expressing the same gene module. (I) EdU pulse-chase analysis of E13.5-E14.5 ActSMO embryos with quantification of %EdU in *Trp53inp1*+ cells (n=3). Cartoon depicts *Trp53inp1*+ cells undergo cell cycle exit immediately following division. Scale bars, 50 µm.

We next applied GT to E13.5 ActSMO dermal scRNA-seq data in which exit and DC differentiation are uncoupled. GT identified four trajectories (Fig. 4D; Supplemental Fig. 3C). Two corresponded to proliferation (S/G2/M) and lower dermal differentiation, whereas two additional trajectories were unique to ActSMO dermis. One contained DC genes and the second lacked DC markers. Cells expressing this latter program localized to a subset of Wnt-active cells and were largely restricted to G1/G0 phase (Fig. 4E). Pathway analysis revealed enrichment for TP53-associated cell-cycle inhibitory genes, including *Trp53inp1* and *Plk2,* overlapping with *Cdkn1a* expression (Fig. 4F; Supplemental Table S1). FISH confirmed significant upregulation of both *Cdkn1a* and *Trp53inp1* in the E13.5 ActSMO dermis relative to controls (Fig. 4G, 3D; Supplemental Fig. S3D)

*Trp53inp1+* ActSMO cells exhibited significantly higher *Lef1* levels than control upper dermal cells (Fig. 4G), consistent with SHH-dependent elevation of Wnt activity and quiescence (Fig. 3D). To test whether elevated Wnt signaling alone induces this program, we applied GT to E13.5 Actβcat dermis, which induces *Cdkn1a* but not DC genes (Supplemental Fig. S3E). A similar *Trp53inp1*-containing trajectory was identified in Actβcat mutants (Supplemental Table S1). Module projection (label transfer) of the ActSMO trajectory onto Actβcat and control datasets revealed a shared molecular state in both mutants that was absent in controls (Fig. 4H).

To determine whether this program reflects a general consequence of reduced proliferation, we analyzed E13.5 Wntless conditional knockout embryos (*K14Cre;Wls^fl/fl^*) in which epidermal Wnt ligand secretion is disrupted and dermal proliferation is reduced (Chen et al. 2012; Fu and Hsu 2013; Qu et al. 2024). Label transfer of the ActSMO/Actβcat-specific trajectory was not detected in Wntless cKO dermis (Fig. 4H; Supplemental Fig. S3F). Notably, dermal proliferation is reduced in Wntless mutants despite the absence of this program, indicating that the Exit trajectory reflects a Wnt-dependent developmental transition rather than generic cell-cycle arrest.

Finally, we examined cell-cycle dynamics of cells expressing this program. At E13.5 and E14.5, most ActSMO *Trp53inp1*+ cells showed low EdU incorporation comparable to quiescent DC cells (Fig. 4I; Supplemental Fig. S4A). EdU pulse-chase experiments showed that many *Trp53inp1*+ cells retained EdU similarly to wildtype DCs (Fig. 4I; Supplemental Fig.S1D) (Gupta et al. 2019; Qu et al. 2022). Although ActSMO dermal clusters contained proliferating Sox2+ cells at E14.5, most became quiescent by E15.5 (Supplemental Fig. S4B), indicating that they exit the cell cycle after division. Together, these data define a shared gene program underlying cell-cycle exit during the pre-DC-to-DC transition, which we term the *“Exit” gene trajectory*.

### Loss of GLI3 chromatin binding induces cell-cycle exit downstream of elevated Wnt signaling

Although Wnt activity is required for dermal proliferation, our data indicate that elevated Wnt activity promotes exit during DC formation (Chen et al. 2012; Gupta et al. 2019). To examine the mechanism, we profiled CTNNB1 (β-catenin) in E13.5 Actβcat and control dermis.

CTNNB1 binding was enriched at canonical Wnt targets (e.g., *Sp5*, *Axin2*, *Lef1*), but not at *Cdkn1a,* Exit GT genes, or other loci associated with cell cycle inhibition (Supplemental Fig. S5A). This raised the question of how elevated Wnt activity promotes exit without direct CTNNB1 binding at inhibitory genes. Because Wnt activity is elevated upon SHH activation, components of the SHH pathway could mediate Wnt-dependent exit. In other tissues, Wnt signaling regulates *Gli3* independently of SHH (Alvarez-Medina et al. 2008). Prior to SHH expression, GLI3 is predominantly present as a cleaved repressor that binds many SHH target loci (te Welscher et al. 2002; Vokes et al. 2008). FISH confirmed *Gli3* expression in Wnt-active dermal cells of control and Actβcat embryos at E13.5 prior to SHH expression (Fig. 5A). To profile GLI3 binding, we performed CUT&RUN in E13.5 Actβcat and control dermis using *Gli3Flag* embryos. GLI3 occupancy was globally reduced in the Actβcat condition reflected by decreased reads at transcriptional start sites (Fig. 5B). De novo motif analysis confirmed enrichment of GLI motifs at regions with reduced binding (Supplemental Fig. S5B). These regions were enriched for genes associated with SHH signaling, although canonical SHH targets (e.g., *Ptch1*, *Gli1*) were not expressed and lacked active histone marks at E13.5 (Fig. 5C). These observations are consistent with recent work showing that GLI3 binding prior to SHH does not actively repress (i.e., inert) many target loci prior to SHH expression (Lex et al. 2022). Western Blot confirmed that cleaved Gli3 protein levels were retained in the Actβcat mutant, indicating that reduced chromatin occupancy was not due to loss of Gli3 protein (Fig. 5D). Thus, elevated Wnt signaling alone reduces GLI3 chromatin binding independently of SHH.

**Fig. 5:**
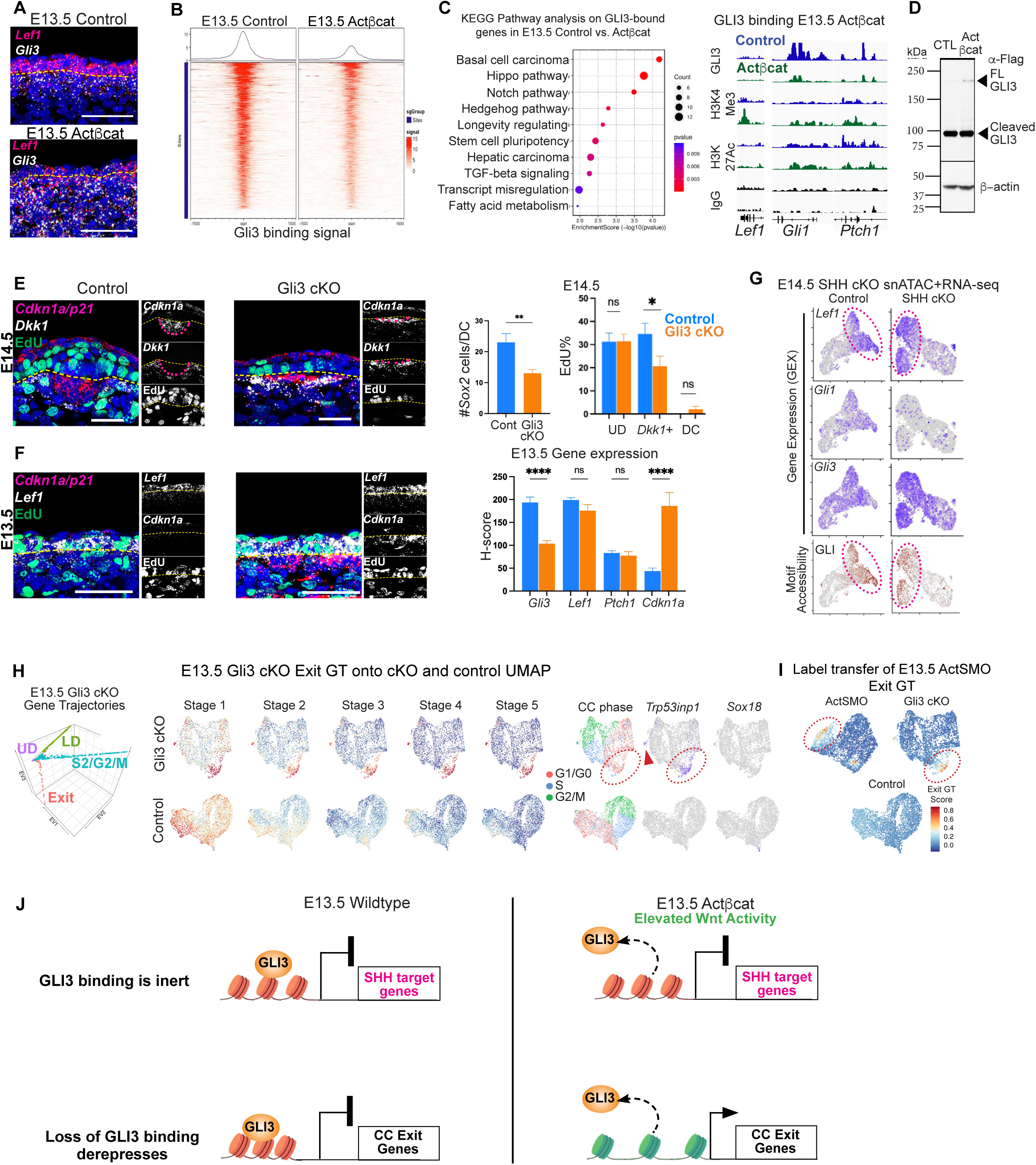
Loss of GLI3 chromatin binding induces premature cell-cycle exit downstream of elevated Wnt signaling. (A) *Gli3* expression in E13.5 control and Actβcat skin prior to SHH. (B) Heat map of GLI3 binding at transcriptional start sites (TSS) in E13.5 control and Actβcat skin. (C) KEGG pathway analysis of GLI3 unbound regions in Actβcat with genome browser tracks at representative loci. (D) Western blot of control and Actβcat lysates probed for Flag (Gli3) and β-actin (n=3). (E) FISH of E14.5 control and Gli3 cKO dermis with quantification of DC size and %EdU by cell type (n=3). (F) FISH of E13.5 control and Gli3 cKO with quantification of transcripts (n=4). (G) Integrated snATAC+RNA-seq UMAPs of E14.5 control and epidermal SHH cKO showing gene expression and GLI motif accessibility. (H) GeneTrajectory diffusion map and UMAPs of E13.5 Gli3 cKO dermal cells, identifying an Exit trajectory. (I) Label transfer of ActSMO Exit trajectory onto Gli3 cKO and control scRNA-seq datasets. (J) Model illustrating that elevated Wnt activity reduces GLI3 chromatin occupancy to derepress genes mediating exit, whereas SHH signaling is required for canonical SHH targets and DC genes. Scale bars, 50 µm.

To test whether loss of GLI3 binding induces exit, we ablated *Gli3* in the dermis (*Axin2CreER;Gli3^fl/fl^* or Gli3 cKO). At E14.5, DCs in Gli3 cKO embryos were smaller than controls, and *Dkk1*+ peri-DC progenitors showed premature *Cdkn1a* upregulation and reduced EdU incorporation (Fig. 5E; Supplemental Fig. S5C). Similarly, at E13.5 before DC formation, Gli3 cKO dermal cells exhibited increased *Cdkn1a* expression and reduced proliferation, phenocopying the Actβcat dermis (Fig. 5F; Supplemental Fig. S5D).

FISH confirmed reduced *Gli3* expression in E13.5 Gli3 cKO dermis, while Wnt targets were unaffected, indicating that GLI3 acts downstream of Wnt activity (Fig. 5F; Supplemental Fig. S5E). Canonical SHH targets remained inactive, showing that loss of GLI3 does not broadly derepress SHH targets. To assess whether GLI3 binding is associated with chromatin organization, we performed chromatin capture (Micro-C) in E13.5 wildtype dermis. GLI3-bound loci were modestly enriched in open chromatin compartments but rarely formed chromatin loops and lacked active promoter marks (Supplemental Fig. S5F). In contrast, canonical Wnt targets (e.g., *Lef1* and *Sp5)* formed chromatin loops and showed active histone modifications. Moreover, snATAC+RNA-seq analysis of E14.5 embryos lacking epidermal *Shh* showed GLI motif accessibility localized to Wnt-active cells even without GLI activator function (Fig. 5G). These results suggest that GLI3 does not broadly repress transcription but may prime regulatory regions prior to SHH activation.

scRNA-seq analysis of E13.5 Gli3 cKO dermis showed reduced *Gli3* expression with retained Wnt target gene expression (Supplemental Fig. S6A,B). GT analysis identified four trajectories, including a trajectory absent in controls that overlapped with the Exit GT identified in the Actβcat and ActSMO mutants (Fig. 5H; Supplemental Fig. S6C). This trajectory was enriched for cell-cycle inhibitory genes, including *Trp53inp1*. Label transfer of the ActSMO Exit GT onto the Gli3 cKO dataset identified a corresponding population that expressed this gene program (Fig. 5I), indicating that this Exit program is shared across ActSMO, Actβcat, and Gli3 cKO conditions.

EdU pulse-chase experiments showed that *Cdkn1a*-high cells in the Gli3 cKO were largely quiescent but represented recent progeny of cell divisions (Supplemental Fig. S5D). Cross-referencing genes that lost GLI3 binding in Actβcat dermis with genes upregulated in Gli3 cKO dermis identified multiple cell-cycle inhibitory genes, including *Mxd4* and *Efna5* (Supplemental Fig. S6D). Exit GT genes such as *Baiap2* and *Cpt1c* also showed reduced GLI3 binding and increased expression. Together, these data indicate that β-catenin and Gli3 act in a shared pathway regulating cell-cycle exit in which elevated Wnt signaling reduces GLI3 chromatin occupancy and induces genes associated with exit (Fig. 5J).

### Uniformly high SHH activity induces dermal condensate gene expression across the Wnt gradient

High SHH activity can induce DC gene expression before cell-cycle exit, indicating that DC differentiation does not strictly require a preceding quiescent state (Fig. 3B). However, in the wildtype condition, DC genes are induced abruptly as cells exit the cell cycle, suggesting that Wnt activity contributes to the timing of DC gene induction. Given that SHH cell-autonomously elevates Wnt activity, we examined whether Wnt signaling levels modulate how DC genes are induced in response to high SHH activity.

To address this, we analyzed the DC gene trajectory in the E13.5 ActSMO mutant, in which SHH activity is uniformly high across cells spanning a range of Wnt activity (Fig. 6A; Supplemental Fig S7A). In the E14.5 wildtype DC gene trajectory, DC genes are induced primarily at the terminal stage, coinciding with the highest levels of *Lef1* and *Ptch1* expression (Fig. 4A,C). By contrast, in the ActSMO condition, DC genes (e.g., *Gal*, *Sox18, Sox2*) were induced progressively across DC trajectory stages, with cumulative expression correlating with increasing *Lef1* levels (Fig. 6B-C). Cells with highest *Lef1* levels showed the greatest progression along the trajectory and the highest DC gene expression. Notably, early stages retained a high proliferative fraction, whereas terminal-stage cells were largely quiescent (G1/G0 fraction ∼0.9). Similarly, ActSMO cells expressing lower levels of the DC marker *Gal* showed high EdU incorporation, whereas *Gal*-high cells were largely non-proliferative like wildtype DC cells (Fig. 6D; Supplemental Fig. S7B).

**Fig. 6:**
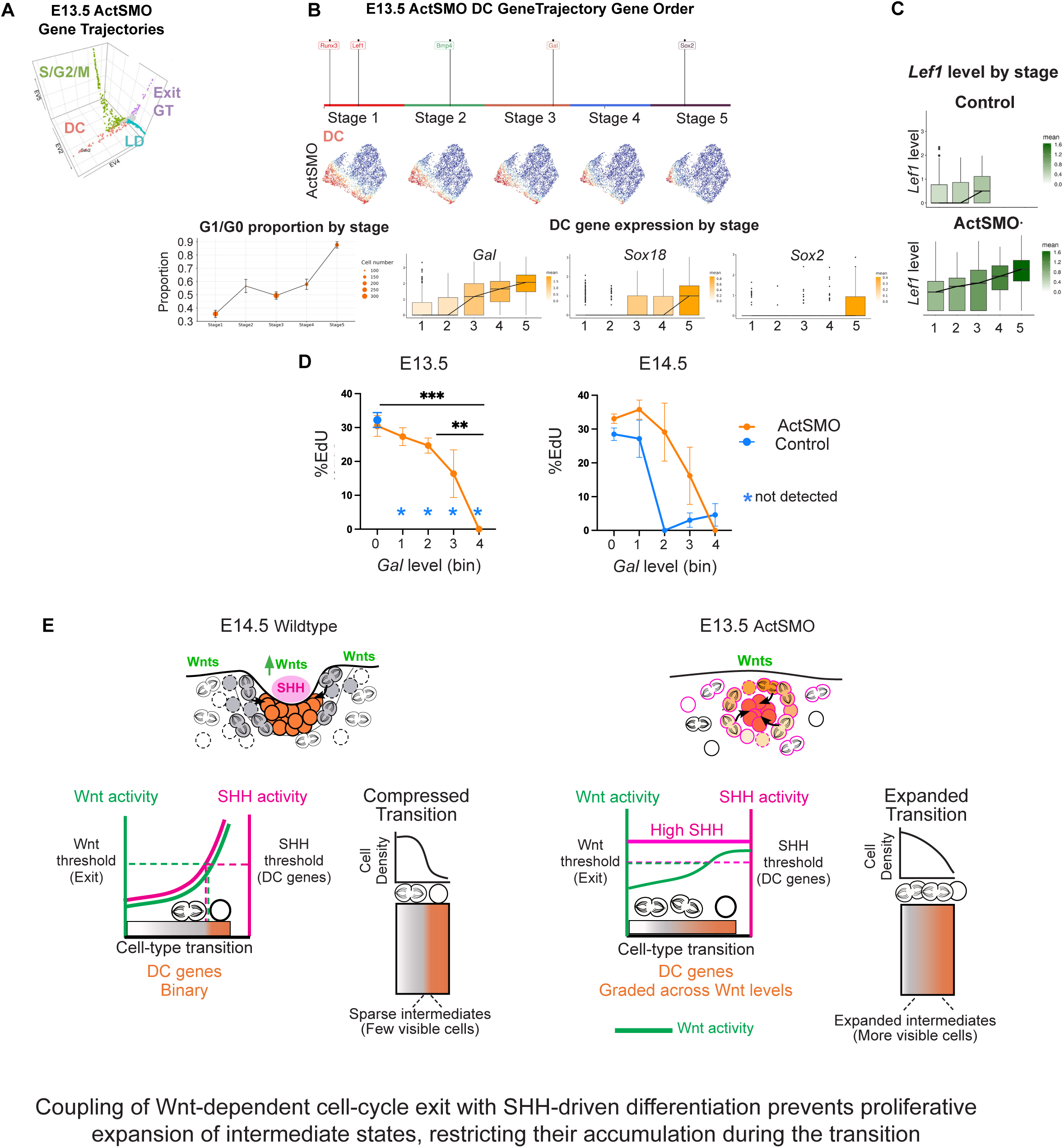
Uniformly high SHH activity induces dermal condensate gene expression across the Wnt gradient. (A) GeneTrajectory diffusion map of E13.5 ActSMO dermal cells. (B) Gene order of the ActSMO DC trajectory with UMAPs colored by trajectory stage and DC gene expression by trajectory stage (bottom). (C) *Lef1* levels by trajectory stage. (D) %EdU by *Gal* level at E13.5 and E14.5 (*, bin level not detected in control). (E) Model illustrating that uniform SHH permits DC gene expression across a broader range of Wnt activity levels, resulting in sequential induction of DC genes correlated with *Lef1* levels. DC genes can be expressed in proliferating cells of lower Wnt activity, expanding intermediate states before coordinated exit. The orange/gray bars represent the extent of intermediate states along the trajectory.

Consistent with the requirement for Wnt signaling in DC differentiation, cells with high SHH activity but lacking Wnt signaling in the ActSMO condition failed to express DC genes (Fig. 6B; Supplemental Fig. S7C). Thus, uniform SHH activation expands the window over which Wnt-competent cells initiate DC gene expression, resulting in graded rather than abrupt induction of DC genes. In both E14.5 wildtype and Actβcat (high Wnt) conditions, DC genes were restricted to Wnt-active cells with high SHH activity, indicating that DC differentiation requires SHH signaling above a defined threshold (Fig. 6D).

### Coupling Wnt and SHH gradients is sufficient to synchronize the timing of cell-cycle exit with molecular differentiation

Because exit and DC differentiation coincide where high Wnt activity overlaps with SHH signaling, we tested whether coupling these gradients is sufficient to restore temporal coordination between the two processes. In wildtype skin, this overlap occurs in dermal cells adjacent to placodes. We therefore induced epidermal SHH expression at E13.5 (*K14Cre;Rosa-LSL-SHH*; SHH OE) to activate SHH signaling in Wnt-active dermal cells before placode formation (Fig. 7A) (Reddy et al. 2001; Phan et al. 2020). FISH analysis showed that *Lef1* and *Ptch1* covaried with dermal depth with highest levels in the most superficial dermis (Fig. 7B). *Lef1* levels were increased relative to E13.5 controls, indicating that SHH enhances Wnt activity. DC genes (e.g., *Sox2*, *Gal*) were restricted superficial layers and coincided with *Cdkn1a* expression and low EdU incorporation (Fig. 7C; Supplemental Fig. S8A). *Dkk1* localized to a deeper dermal layer beneath DC gene-expressing cells. scRNA-seq analysis of E13.5 SHH OE dermis confirmed overlapping *Lef1* and *Ptch1* expression, with DC genes and *Cdkn1a* restricted to cells with the highest combined signal levels (Fig. 7D).

**Fig. 7:**
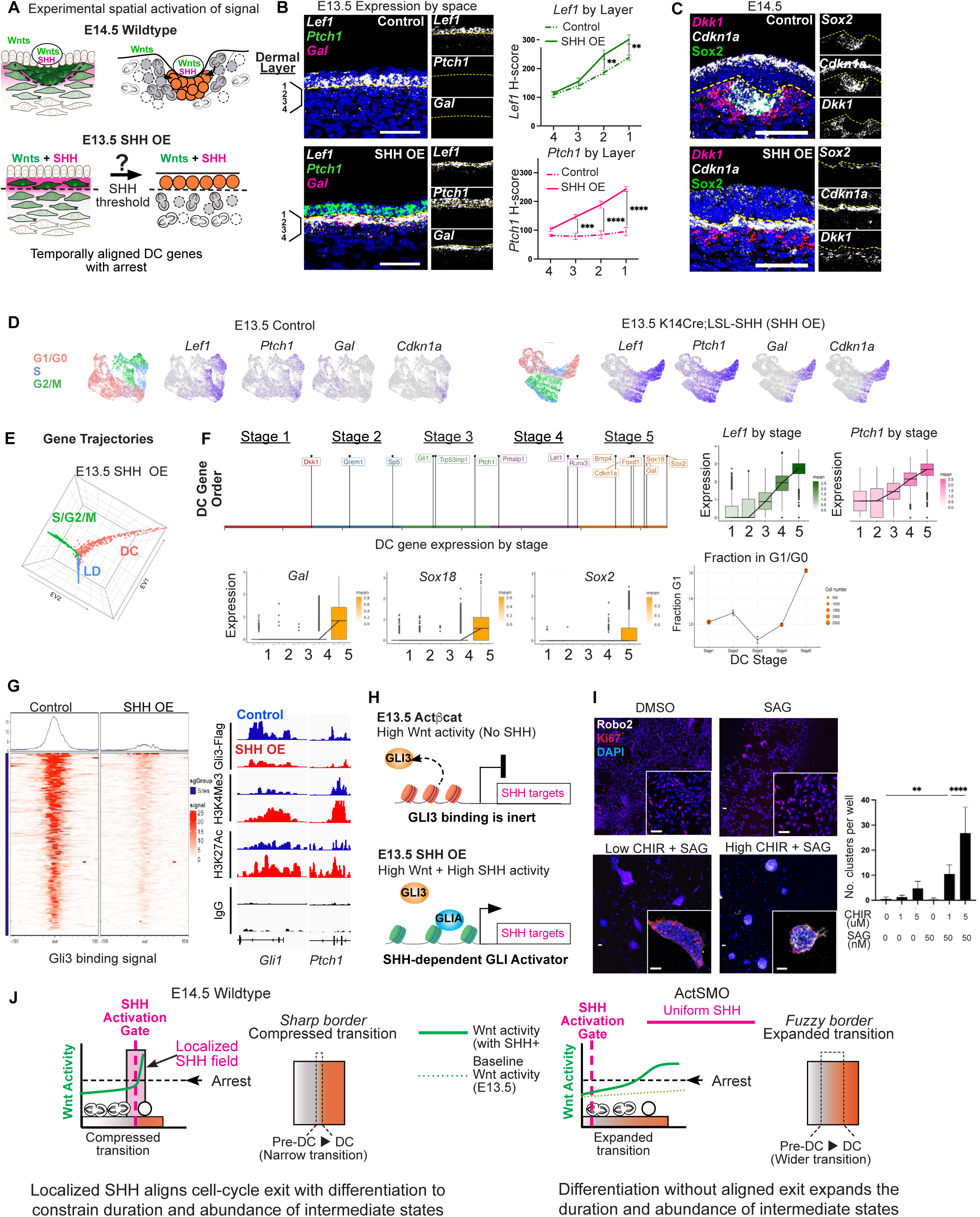
Coordinated Wnt and SHH signaling synchronizes cell-cycle exit and dermal condensate differentiation. (A) Experimental design to test if Wnt and SHH co-activation aligns cell-cycle exit with DC gene expression by epidermal co-expression of Wnt and SHH ligands at E13.5 (SHH OE). (B) FISH of E13.5 control and SHH OE skin showing covarying *Ptch1* and *Lef1* levels across dermal layers with elevated *Lef1* where *Ptch1* is concurrently activated (n=3). (C) E14.5 control and SHH OE skin showing spatial partitioning of cells by cell-type markers with *Cdkn1a* coinciding with *Sox2* expression (n=3). (D) UMAPs of E13.5 control and SHH OE dermal scRNA-seq colored by indicated genes or cell-cycle phase. (E) GeneTrajectory diffusion map of E13.5 SHH OE dermal cells. (F) SHH OE DC trajectory showing DC gene expression and G1/G0 fraction by stage; right, *Lef1* and *Ptch1* levels by stage. (G) Heat map of GLI3 binding at transcriptional start sites (TSS) in E13.5 control and SHH OE skin with genome browser tracks at *Gli1* and *Ptch1*. (H) Reduced GLI3 chromatin binding upon Wnt and SHH coactivation. (I) 3D cultures stained for Robo2 and Ki67 with quantification of dermal clusters. (J) Model illustrating how Wnt and SHH signaling generate sharp boundaries by constraining the duration and abundance of intermediate states during the pre-DC-to-DC transition. In wildtype skin, a localized SHH field aligns cell-cycle exit with differentiation in Wnt-primed cells, limiting the persistence and expansion of intermediate states, producing a sharp boundary. When SHH signaling is uniform (ActSMO), this spatial restriction is lost, allowing differentiation across a broader range of Wnt activity; intermediate states persist and expand, producing a diffuse transition. The pink bar indicates the spatial SHH field and the green dotted line the pre-existing Wnt gradient at E13.5. Scale bars, 50 µm.

GT analysis identified three trajectories corresponding to proliferation, lower dermal differentiation and DC differentiation, similar to E14.5 wildtype dermis (Fig. 4A, 7E; Supplemental Fig. S8B). A distinct Exit trajectory was not detected. Within the DC trajectory, *Lef1* and *Ptch1* increased coordinately, and DC genes and *Cdkn1a* were induced synchronously at the terminal stage, coincident with quiescence (Fig. 7F; Supplemental Fig. S8C). Thus, coupling Wnt and SHH activity restores temporal alignment of differentiation and exit.

We next examined whether GLI3 chromatin occupancy is similarly reduced when Wnt and SHH signaling are coupled. GLI3 CUT&RUN analysis in SHH OE dermis showed global depletion of GLI3 binding (Fig. 7G; Supplemental Fig. S8D), indicating that GLI3 displacement is not an artifact of genetic stabilization of β-catenin. Loss of GLI3 binding occurred at cell cycle regulatory genes (*Mxd4* and *Efna5*) associated with active histone marks (Supplemental Fig. S8E). In contrast to Actβcat mutants, SHH target genes (*Ptch1*, *Gli1*) were transcriptionally activated, accompanied by increased H3K4Me3 and H3K27Ac signal. Because SHH signaling converts cleaved GLI3 repressor into full-length GLI activator, the observed loss of GLI3 occupancy likely reflects replacement by other GLI activators at SHH target loci (Fig. 7H). These data indicate that GLI3 eviction both derepresses exit genes and may prime SHH-dependent target loci for activation.

To test whether graded Wnt and SHH inputs are sufficient to coordinate these processes outside of the tissue context, we reconstituted the system *in vitro* by using E13.5 PDGFRαH2BGFP dermal cells (Hamilton et al. 2003). These cells showed dose-dependent gene responses to the Wnt agonist (CHIR99021) or the Smoothened agonist (SAG) (Supplemental Fig. S9A). SAG enhanced *Lef1* expression in CHIR-treated cells (Supplemental Fig. S9B). High CHIR alone induced *Cdkn1a*, whereas combined CHIR and SAG treatment induced DC genes, including *Gal* and *Sox18*.

To assess morphogenesis, E13.5 dermal cells were seeded onto a collagen-based 3D matrix with varying CHIR and SAG concentrations (Fig. 7I; Supplemental Fig. S9C). High CHIR (5 μM) with SAG (50 nM) produced large dermal clusters with low proliferation that expressed Robo2, a DC gene that regulates cell migration (Goncalves et al. 2020; Guzman-Palma et al. 2021). Clusters from other conditions did not express Robo2 and contained more Ki67+ cells. Thus, coupled Wnt and SHH activity aligns molecular differentiation with exit *in vitro*. Together, these results indicate that the pre-DC–to-DC transition comprises separable biological processes whose timing can be coordinated by Wnt and SHH (Fig. 7J).

## Discussion

In this study, we identify a mechanism by which interacting morphogen signals generate sharp cell-type boundaries by coordinating the timing of separable components of a cell-state transition. During dermal condensate (DC) formation, we show that Sonic Hedgehog (SHH) acts on Wnt-primed cells to elevate Wnt activity, which is sufficient to induce cell-cycle exit through eviction of the transcriptional repressor GLI3 from chromatin, while SHH concurrently initiates the differentiation program. By aligning differentiation with cell-cycle exit, this coordination limits the duration of intermediate states and prevents their proliferative accumulation, causing transitional cells to remain sparse and difficult to detect. Boundary sharpness therefore reflects a compression of a developmental transition in both the duration and abundance of intermediate states, rather than their absence. While sharp transitions can also arise from nonlinear ligand diffusion, our findings reveal a distinct mechanism in which morphogen interactions synchronize concurrent gene programs.

These findings revisit classic interpretations of the French flag problem, which highlight fixed morphogen thresholds as primary determinants of spatial patterning (Driever and Nusslein-Volhard 1988; Roelink et al. 1995; Greenfeld et al. 2021). Our data instead indicate that during DC formation, morphogen thresholds act as triggers that initiate a committed developmental progression. Here, SHH gates entry into the DC differentiation program, whereas Wnt activity regulates the timing and progression of its execution, establishing a division of labor between these pathways. Abrupt boundaries may therefore emerge not from static positional decoding, but from progression through the same developmental trajectory (Fig. 1A) in which aligning differentiation with cell-cycle exit compresses the duration and abundance of intermediate states. Accordingly, early DC states in the ActSMO condition remained proliferative, whereas terminal states were largely quiescent. As a result, intermediate states become more abundant because differentiation occurs before cell-cycle exit, allowing partially differentiated cells to expand.

Although tight coupling of Wnt and SHH activities produces a sharp transition during DC formation, variation in the timing, strength, or spatial alignment of interacting morphogen gradients may support alternative modes of tissue organization. In systems containing transit-amplifying populations, differentiation can proceed without immediate cell-cycle exit, expanding intermediate states and producing graded boundaries (Morita et al. 2021; Sun et al. 2023). Thus, the duration and abundance of transitional states are tunable outputs of morphogen coordination that regulate boundary sharpness across tissues (Fig. 7J). Additional pathways, including FGF signaling, have been implicated in DC formation and may act upstream of SHH in this process (Biggs et al. 2018; Mok et al. 2019; Qu et al. 2022). Broadly, these findings suggest that precise tissue organization in regenerative and engineered systems depends not only on establishing cell identity, but on controlling the duration and abundance of intermediate states during developmental transitions.

How SHH signaling elevates Wnt activity is unlikely to involve direct transcriptional activation of Wnt target genes by GLI factors. Instead, elevated Wnt activity is associated with loss of GLI3 chromatin occupancy at loci involved in cell-cycle regulation. Although GLI3 eviction does not produce large-scale chromatin reorganization, it coincides with SHH-dependent activation of differentiation genes, suggesting that chromatin-level cross-regulation between Wnt and SHH pathways aligns exit with differentiation.

At the tissue level, this SHH-dependent enhancement of Wnt responsiveness shapes how transitions unfold in space and time. Prior to SHH signaling, dermal cells occupy positions along a Wnt-defined continuum, reflecting differences in transcriptional state. These differences encode competence - how rapidly cells traverse the transition once SHH is engaged, rather than pre-specified fates. Cells with higher Wnt activity reach exit more rapidly upon SHH activation, whereas cells with lower Wnt activity transition more slowly, expanding intermediate states. Therefore, SHH establishes a shared transition window across a pre-patterned landscape, compressing the duration of intermediate states locally and producing sharp boundaries without altering the sequence of states traversed. It will be important to determine whether Wnt signaling also contributes to dermal cell migration or aggregation during DC formation.

This work establishes an experimentally tractable mammalian system for investigating how interacting morphogen gradients regulate the timing of concurrent biological processes. Integrating *in vivo* genetics with trajectory-based analysis resolves overlapping differentiation programs that are otherwise difficult to detect due to their limited duration and abundance, revealing how dynamic signaling interactions shape discrete cell-state transitions (Qu et al. 2024).

## Materials and Methods

### Mouse strains and tamoxifen induction

Embryos were staged as days post coitum by vaginal plug detection. Pregnant dams received a single tamoxifen dose (20mg/ml in corn oil, Sigma; 30-60 µg/gm body weight) by gavage at E11.5. *Axin2CreER* (van Amerongen et al. 2012) mice were crossed with *Rosa-LSLSmoM2YFP* (Jeong et al. 2004), *β-catenin^fl(Ex3/+^* (Harada et al. 1999) and *Gli3^fl/fl^* (Blaess et al. 2008) mice. *K14Cre* (Dassule et al. 2000) mice were bred to *Shh* Overexpressor (Wang et al. 2010) and *Wntless^fl/^*^fl^ (Carpenter et al. 2010) mice. *Rosa-Flag-Gli3YFP* (Vokes et al. 2008) mice were crossed with *Axin2CreER* and *β-catenin^fl(Ex3/+)^*mice. *R26Fucci2aR* (Abe et al. 2013) mice were used to visualize cell-cycle phase. *PDGFRaH2BGFP* (Hamilton et al. 2003) mice were used for dermal cultures. A random population of both male and female mice were used. All procedures involving animal subjects were performed under the approval of the Institutional Animal Care and Use Committee of the Yale School of Medicine.

### EdU Incorporation Assay

EdU (25 µg/gm) was administered to pregnant mice intraperitoneally, and embryos were harvested after indicated hours. EdU was detected using the Click-it EdU Imaging kit (Alexa 555 or Alexa 488; Life technologies) following manufacturer’s instructions.

### Whole mount immunofluorescence

Dorsolateral skin was microdissected, placed on nucleopore filters (VWR) and fixed with 4% PFA. Explants were blocked and incubated overnight with anti-Sox2 (1:400, Abcam), anti-GFP (1:500, Abcam), anti-RFP (1:500, Rockland) antibodies. Explants were washed and incubated overnight with Alexa Fluor 488 or 568 anti-rabbit or anti-chicken (1:200, Invitrogen) antibodies. Explants were washed and nuclear counterstained (Hoechst) before mounting onto slides.

### Fluorescent In-situ hybridization

Embryos were fixed in 10% formalin, paraffin embedded and sectioned. RNA FISH was performed using the RNAscope Multiplex Fluorescent Detection Kit (ACDBio) according to manufacturer’s instructions. Probe detection was done using HRP-C1-3 reagents and tyramide signal amplification (VIVID 650 and 570). EdU staining was done as described above. Probes used: Mm-Lef1 (441861), Mm-Ptch1 (402811), Mm-Dkk1 (402521), Mm-Sox2 (401041), Mm-Gal (400961), Mm-Gli3 (400961) and Mm-Trp53inp1 (1161531).

### Microscopy

FISH images were acquired using the Leica Stellaris 8 DMi8 confocal microscope with a 40X oil immersion objective lens, scanned at 5 µm thickness, 1024 × 1024-pixel width. Skin explants were imaged using the LaVision TriM Scope II (LaVision Biotec) microscope equipped with a Chameleon Vision II (Coherent) two-photon laser (810 -1000 nm) for 3-D images ranging from 50-120 µm (2 µm serial optical sections) using a 20X water immersion lens (Olympus), scanned with a field of view of 0.3–0.5 mm^2^ at 800 Hz.

### Single-cell dissociation

Embryonic dorsolateral skin was microdissected and pooled after genotyping (2-3 embryos per condition) and dissociated into a single-cell suspension in 0.25% trypsin (Life Technologies) for 15 minutes at 37°C. DAPI-excluded live skin cells were FACS sorted and submitted for 10X Genomics library preparation at 1.0×10^6^/mL or processed for CUT&RUN or micro-C experiments.

### Single-cell RNA sequencing

Single-cell libraries were generated using the 10X Chromium Single Cell 3′ v2 platform and sequenced on an Illumina NovaSeq 6000. Raw reads were processed using the Cell Ranger pipeline and aligned to the mm10 genome to generate gene-cell count matrices. Downstream analysis was performed in Seurat. Cells with >1,000 detected genes, <25% mitochondrial content, and >5% ribosomal content were retained, and doublets were removed using scDblFinder. Data were log-normalized, and principal component analysis was performed on the top variable genes. UMAP was used for visualization and clustering was performed using Seurat FindClusters (resolution 0.5). Clusters expressing *Col1a1* were classified as dermal, and clusters expressing *Lef1* or *Dkk1* were defined as upper dermal.

### GeneTrajectory inference

GeneTrajectory analysis was performed as described (Qu et al. 2024). Highly variable genes were used to construct diffusion maps based on the top 20 PCs. Cell k-nearest neighbor graphs (k = 10) were generated from the top diffusion components. Gene trajectories and time steps were determined by inspection of gene embeddings. Each trajectory was partitioned into 5–6 gene bins, and cells were assigned to stages based on expression of ≥50% of genes in the corresponding bin. Exit gene trajectory scores were calculated using Seurat AddModuleScore based on genes identified in the E13.5 ActSMO dataset. Pathway enrichment analysis was performed using ReactomePA, and cell-cycle phase was estimated using Seurat cell-cycle scoring. For comparative analysis, upper dermal cells from paired conditions were merged and analyzed using standard Seurat workflows. Differential abundance was assessed using DA-seq, and differential gene expression was performed using Seurat FindMarkers.

### Integrated snATAC+RNA-seq Multiomic Analysis

snRNA-seq and snATAC-seq data from E14.5 *K14Cre;SHH^fl/fl^* and control skin were processed using Seurat and Signac. RNA data were normalized using SCTransform, and ATAC data were reduced using latent semantic indexing (LSI). Modalities were integrated using weighted nearest neighbor (WNN) analysis, and joint UMAP embeddings were generated for visualization. Transcription factor motif accessibility was calculated using chromVAR.

### CUT&RUN experiments

CUT&RUN was done using the CUTANA ChIC/CUT&RUN kit (Epicypher) following the manufacturer’s protocol. ∼5×10⁵ cells were used for nuclei isolation and immobilization on ConA beads. Nuclei were incubated with primary antibodies (0.5 µg/reaction): IgG, H3K27Ac, H3K4me3, FLAG, and CTNNB1. Libraries were prepared using the CUTANA library prep kit and sequenced on an Illumina NovaSeq 6000. Reads were quality filtered, trimmed, and aligned to the mm9 genome using Bowtie2. BAM files were processed using SAMtools, and peaks were called with MACS2 using IgG as control. Coverage tracks were generated using deepTools and visualized in IGV. Differential binding analysis was perfumed using DESeq2. Heatmaps were generated using DiffBind and motif enrichment analysis was performed using HOMER.

### Micro-C Experiments

Micro-C was performed using the Dovetail Micro-C kit (Cantata Bio) following the manufacturer’s protocol. E13.5 dissociate skin cells were crosslinked with formaldehyde. Chromatin was digested with micrococcal nuclease, proximity-ligated, reverse crosslinked, and purified. Libraries were prepared using the Dovetail library module and sequenced to 300 million reads per biological replicate. Reads were aligned to the mm10 genome using BWA-MEM. Valid ligation pairs were identified and filtered using the Pairtools pipeline. Contact matrices were generated at 5-kb resolution using cooler and normalized by iterative correction. Chromatin loops were identified using Mustache and loop anchors intersected with CUT&RUN peaks using Bedtools. A/B compartments were calculated at 40-kb resolution using FAN-C. Interaction maps and compartment tracks were visualized using FAN-C.

### Western blot

Cells were lysed in RIPA buffer, and proteins from clarified lysates were resolved on SDS–PAGE gels and transferred to PVDF membranes. Membranes were blocked in 5% BSA/TBST, incubated with primary antibodies (anti-FLAG and anti-Actin) overnight at 4°C, washed, and incubated with HRP-conjugated secondary antibodies. Proteins were detected by chemiluminescence.

### Dermal cultures

E13.5 PDGFRαH2BGFP dermal cells were FACS-isolated. GFP⁺ cells were cultured in DMEM with 10% FBS, penicillin–streptomycin, L-glutamine, and plated at 3×10⁶ cells per well in 6-well plates. For chemical perturbation assays, cells were treated with CHIR99021 or SAG as indicated for 48–72 h. For 3D cultures, 3×10⁵ cells were seeded in µ-slide 8-well glass-bottom chambers (Ibidi) precoated with 1.5 mg/mL polymerized collagen type I (Corning) before seeding. Cultures were fixed in 4% PFA and blocked in 5% normal donkey serum, 1% BSA, 0.2% Triton X-100 and incubated overnight at 4°C with rat anti-Ki67 (Invitrogen) and rabbit anti-Robo2 (Cell Signaling), followed by Alexa Fluor–conjugated secondary antibodies (Invitrogen) and nuclear counterstain before imaging by confocal microscopy.

### Quantitative PCR

RNA was isolated using the RNeasy Mini Kit (Qiagen) and reverse-transcribed using SuperScript III (Invitrogen). QPCR was performed using SYBR Green Master Mix (Thermo Fisher) on a Viia7 real-time PCR system. Reactions were run in technical triplicates and normalized to *Gapdh*. Relative expression was calculated using the ΔΔCt method.

### Image analysis and quantification

Images were analyzed using ImageJ and Adobe Photoshop. scRNA-seq visualizations were generated using ggplot2 in R. DC cell counts were quantified from 3D whole-mount mosaics (1000×1000 µm FOV; 60–100 µm depth); n=3–4 embryos. Sox2⁺, EdU⁺ cells and RFP⁺ cells were manually counted in ImageJ using orthogonal views. Upper dermis was defined as 10–20 µm below the epidermis. Peri-DC was defined as cells within two cell layers of Sox2⁺ DCs. For RNA FISH, cells with 4–5 dots were considered positive and expression levels were quantified by H-score based on weighted dot counts per cell.

### Statistical analysis

All statistical values are expressed as mean ± SEM. Two-group comparisons used unpaired Student’s t-test; multiple comparisons used one-way ANOVA with Tukey’s test (\**P*<0.05, \*\**P*<0.01, \*\*\**P*<0.001, \*\*\*\**P*<0.0001).

## Data and code availability

Genomic datasets will be deposited in the NCBI Gene Expression Omnibus and made publicly available upon publication.

## Acknowledgements

SHH overexpressor mice were provided by David Wang. This work was supported by National Institutes of Health grants R01AR076420 (P.M.); R01GM131642, UM1PA051410, U54AG076043, U54AG079759, U01DA053628, P50CA121974, R33DA047037 (Y.K.); LEO Foundation LF-OC-23-001347 (P.M.),

## Author Contributions

Conceptualization, P.M., Y.K., R.L., Y.J., S.P.; Methodology, P.M., R.L., Y.K., Y.J., S.P., E.B., S.V., H.L.; Software, R.L., Y.K.; Resources, M.T., K.P.; Investigation, P.M., R.L., Y.J., S.P. H.L, E.B., S.V.; Writing – Original Draft, P.M.; Writing – Review & Editing, P.M., R.L., Y.J., S.P., T.X., R.D., Y.K., C.L., S.L., E.B.; Supervision, P.M., Y.K.; Funding Acquisition, P.M.

## References

Abe T, Sakaue-Sawano A, Kiyonari H, Shioi G, Inoue K, Horiuchi T, Nakao K, Miyawaki A, Aizawa S, Fujimori T. 2013. Visualization of cell cycle in mouse embryos with Fucci2 reporter directed by Rosa26 promoter. Development 140: 237–246.

Alvarez-Medina R, Cayuso J, Okubo T, Takada S, Marti E. 2008. Wnt canonical pathway restricts graded Shh/Gli patterning activity through the regulation of Gli3 expression. Development 135: 237–247.

Benzinger D, Briscoe J. 2025. Investigating morphogen and patterning dynamics with optogenetic control of morphogen production. Dev Cell 60: 3421–3430 e3426.

Biggs LC, Makela OJ, Myllymaki SM, Das Roy R, Narhi K, Pispa J, Mustonen T, Mikkola ML. 2018. Hair follicle dermal condensation forms via Fgf20 primed cell cycle exit, cell motility, and aggregation. Elife 7.

Blaess S, Stephen D, Joyner AL. 2008. Gli3 coordinates three-dimensional patterning and growth of the tectum and cerebellum by integrating Shh and Fgf8 signaling. Development 135: 2093–2103.

Carpenter AC, Rao S, Wells JM, Campbell K, Lang RA. 2010. Generation of mice with a conditional null allele for Wntless. Genesis 48: 554–558.

Chase HB, Rauch R, Smith VW. 1951. Critical stages of hair development and pigmentation in the mouse. Physiol Zool 24: 1–8.

Chen D, Jarrell A, Guo C, Lang R, Atit R. 2012. Dermal beta-catenin activity in response to epidermal Wnt ligands is required for fibroblast proliferation and hair follicle initiation. Development 139: 1522–1533.

Chiang C, Swan RZ, Grachtchouk M, Bolinger M, Litingtung Y, Robertson EK, Cooper MK, Gaffield W, Westphal H, Beachy PA et al. 1999. Essential role for Sonic hedgehog during hair follicle morphogenesis. Dev Biol 205: 1–9.

Chuong CM, Widelitz RB, Ting-Berreth S, Jiang TX. 1996. Early events during avian skin appendage regeneration: dependence on epithelial-mesenchymal interaction and order of molecular reappearance. J Invest Dermatol 107: 639–646.

Dassule HR, Lewis P, Bei M, Maas R, McMahon AP. 2000. Sonic hedgehog regulates growth and morphogenesis of the tooth. Development 127: 4775–4785.

Driever W, Nusslein-Volhard C. 1988. The bicoid protein determines position in the Drosophila embryo in a concentration-dependent manner. Cell 54: 95–104.

Fu J, Hsu W. 2013. Epidermal Wnt controls hair follicle induction by orchestrating dynamic signaling crosstalk between the epidermis and dermis. J Invest Dermatol 133: 890–898.

Glover JD, Wells KL, Matthaus F, Painter KJ, Ho W, Riddell J, Johansson JA, Ford MJ, Jahoda CAB, Klika V et al. 2017. Hierarchical patterning modes orchestrate hair follicle morphogenesis. PLoS Biol 15: e2002117.

Goncalves AN, Correia-Pinto J, Nogueira-Silva C. 2020. ROBO2 signaling in lung development regulates SOX2/SOX9 balance, branching morphogenesis and is dysregulated in nitrofen-induced congenital diaphragmatic hernia. Respir Res 21: 302.

Greenfeld H, Lin J, Mullins MC. 2021. The BMP signaling gradient is interpreted through concentration thresholds in dorsal-ventral axial patterning. PLoS Biol 19: e3001059.

Gregor T, Wieschaus EF, McGregor AP, Bialek W, Tank DW. 2007. Stability and nuclear dynamics of the bicoid morphogen gradient. Cell 130: 141–152.

Gritli-Linde A, Hallberg K, Harfe BD, Reyahi A, Kannius-Janson M, Nilsson J, Cobourne MT, Sharpe PT, McMahon AP, Linde A. 2007. Abnormal hair development and apparent follicular transformation to mammary gland in the absence of hedgehog signaling. Dev Cell 12: 99–112.

Gupta K, Levinsohn J, Linderman G, Chen D, Sun TY, Dong D, Taketo MM, Bosenberg M, Kluger Y, Choate K et al. 2019. Single-Cell Analysis Reveals a Hair Follicle Dermal Niche Molecular Differentiation Trajectory that Begins Prior to Morphogenesis. Dev Cell 48: 17–31 e16.

Guzman-Palma P, Contreras EG, Mora N, Smith M, Gonzalez-Ramirez MC, Campusano JM, Sierralta J, Hassan BA, Oliva C. 2021. Slit/Robo Signaling Regulates Multiple Stages of the Development of the Drosophila Motion Detection System. Front Cell Dev Biol 9: 612645.

Hamilton TG, Klinghoffer RA, Corrin PD, Soriano P. 2003. Evolutionary divergence of platelet-derived growth factor alpha receptor signaling mechanisms. Mol Cell Biol 23: 4013–4025.

Harada N, Tamai Y, Ishikawa T, Sauer B, Takaku K, Oshima M, Taketo MM. 1999. Intestinal polyposis in mice with a dominant stable mutation of the beta-catenin gene. EMBO J 18: 5931–5942.

Hardy MH. 1992. The secret life of the hair follicle. Trends Genet 8: 55–61.

Jeong J, Mao J, Tenzen T, Kottmann AH, McMahon AP. 2004. Hedgehog signaling in the neural crest cells regulates the patterning and growth of facial primordia. Genes Dev 18: 937–951.

Kicheva A, Bollenbach T, Wartlick O, Julicher F, Gonzalez-Gaitan M. 2012. Investigating the principles of morphogen gradient formation: from tissues to cells. Curr Opin Genet Dev 22: 527–532.

Lander AD, Nie Q, Wan FY. 2002. Do morphogen gradients arise by diffusion? Dev Cell 2: 785–796.

Lex RK, Zhou W, Ji Z, Falkenstein KN, Schuler KE, Windsor KE, Kim JD, Ji H, Vokes SA. 2022. GLI transcriptional repression is inert prior to Hedgehog pathway activation. Nat Commun 13: 808.

Millar SE. 2002. Molecular mechanisms regulating hair follicle development. J Invest Dermatol 118: 216–225.

Mok KW, Saxena N, Heitman N, Grisanti L, Srivastava D, Muraro MJ, Jacob T, Sennett R, Wang Z, Su Y et al. 2019. Dermal Condensate Niche Fate Specification Occurs Prior to Formation and Is Placode Progenitor Dependent. Dev Cell 48: 32–48 e35.

Morita R, Sanzen N, Sasaki H, Hayashi T, Umeda M, Yoshimura M, Yamamoto T, Shibata T, Abe T, Kiyonari H et al. 2021. Tracing the origin of hair follicle stem cells. Nature 594: 547–552.

Noramly S, Freeman A, Morgan BA. 1999. beta-catenin signaling can initiate feather bud development. Development 126: 3509–3521.

Ouspenskaia T, Matos I, Mertz AF, Fiore VF, Fuchs E. 2016. WNT-SHH Antagonism Specifies and Expands Stem Cells prior to Niche Formation. Cell 164: 156–169.

Phan QM, Fine GM, Salz L, Herrera GG, Wildman B, Driskell IM, Driskell RR. 2020. Lef1 expression in fibroblasts maintains developmental potential in adult skin to regenerate wounds. Elife 9.

Qu R, Cheng X, Sefik E, Stanley Iii JS, Landa B, Strino F, Platt S, Garritano J, Odell ID, Coifman R et al. 2024. Gene trajectory inference for single-cell data by optimal transport metrics. Nat Biotechnol.

Qu R, Gupta K, Dong D, Jiang Y, Landa B, Saez C, Strickland G, Levinsohn J, Weng PL, Taketo MM et al. 2022. Decomposing a deterministic path to mesenchymal niche formation by two intersecting morphogen gradients. Dev Cell 57: 1053–1067 e1055.

Reddy S, Andl T, Bagasra A, Lu MM, Epstein DJ, Morrisey EE, Millar SE. 2001. Characterization of Wnt gene expression in developing and postnatal hair follicles and identification of Wnt5a as a target of Sonic hedgehog in hair follicle morphogenesis. Mech Dev 107: 69–82.

Roelink H, Porter JA, Chiang C, Tanabe Y, Chang DT, Beachy PA, Jessell TM. 1995. Floor plate and motor neuron induction by different concentrations of the amino-terminal cleavage product of sonic hedgehog autoproteolysis. Cell 81: 445–455.

Rompolas P, Deschene ER, Zito G, Gonzalez DG, Saotome I, Haberman AM, Greco V. 2012. Live imaging of stem cell and progeny behaviour in physiological hair-follicle regeneration. Nature 487: 496–499.

St-Jacques B, Dassule HR, Karavanova I, Botchkarev VA, Li J, Danielian PS, McMahon JA, Lewis PM, Paus R, McMahon AP. 1998. Sonic hedgehog signaling is essential for hair development. Curr Biol 8: 1058–1068.

Sun Q, Lee W, Hu H, Ogawa T, De Leon S, Katehis I, Lim CH, Takeo M, Cammer M, Taketo MM et al. 2023. Dedifferentiation maintains melanocyte stem cells in a dynamic niche. Nature 616: 774–782.

te Welscher P, Zuniga A, Kuijper S, Drenth T, Goedemans HJ, Meijlink F, Zeller R. 2002. Progression of vertebrate limb development through SHH-mediated counteraction of GLI3. Science 298: 827–830.

van Amerongen R, Bowman AN, Nusse R. 2012. Developmental stage and time dictate the fate of Wnt/beta-catenin-responsive stem cells in the mammary gland. Cell Stem Cell 11: 387–400.

Vokes SA, Ji H, Wong WH, McMahon AP. 2008. A genome-scale analysis of the cis-regulatory circuitry underlying sonic hedgehog-mediated patterning of the mammalian limb. Genes Dev 22: 2651–2663.

Wolpert L. 1969. Positional information and the spatial pattern of cellular differentiation. J Theor Biol 25: 1–47.

Woo WM, Zhen HH, Oro AE. 2012. Shh maintains dermal papilla identity and hair morphogenesis via a Noggin-Shh regulatory loop. Genes Dev 26: 1235–1246.

Zhang Y, Tomann P, Andl T, Gallant NM, Huelsken J, Jerchow B, Birchmeier W, Paus R, Piccolo S, Mikkola ML et al. 2009. Reciprocal requirements for EDA/EDAR/NF-kappaB and Wnt/beta-catenin signaling pathways in hair follicle induction. Dev Cell 17: 49–61.

